# Robust Benchmark Structural Variant Calls of An Asian Using the State-of-Art Long Fragment Sequencing Technologies

**DOI:** 10.1101/2020.08.10.245308

**Authors:** Xiao Du, Lili Li, Fan Liang, Sanyang Liu, Wenxin Zhang, Shuai Sun, Yuhui Sun, Fei Fan, Linying Wang, Xinming Liang, Weijin Qiu, Guangyi Fan, Ou Wang, Weifei Yang, Jiezhong Zhang, Yuhui Xiao, Yang Wang, Depeng Wang, Shoufang Qu, Fang Chen, Jie Huang

## Abstract

The importance of structural variants (SVs) on phenotypes and human diseases is now recognized. Although a variety of SV detection platforms and strategies that vary in sensitivity and specificity have been developed, few benchmarking procedures are available to confidently assess their performances in biological and clinical research. To facilitate the validation and application of those approaches, our work established an Asian reference material comprising identified benchmark regions and high-confidence SV calls. We established a high-confidence SV callset with 8,938 SVs in an EBV immortalized B lymphocyte line, by integrating four alignment-based SV callers [from 109× PacBio continuous long read (CLR), 22× PacBio circular consensus sequencing (CCS) reads, 104× Oxford Nanopore long reads, and 114× optical mapping platform (Bionano)] and one *de novo* assembly-based SV caller using CCS reads. A total of 544 randomly selected SVs were validated by PCR and Sanger sequencing, proofing the robustness of our SV calls. Combining trio-binning based haplotype assemblies, we established an SV benchmark for identification of false negatives and false positives by constructing the continuous high confident regions (CHCRs), which cover 1.46Gb and 6,882 SVs supported by at least one diploid haplotype assembly. Establishing high-confidence SV calls for a benchmark sample that has been characterized by multiple technologies provides a valuable resource for investigating SVs in human biology, disease, and clinical diagnosis.

## Introduction

SVs are generally defined as genomic changes spanning at least 50 base pairs (bp), including deletions, insertions, duplications, inversions, and translocations [1]. They contribute to the diversity and evolution of human genomes at individual and population levels [2, 3]. Owing to their large size, SVs often exert greater impacts on gene functions and phenotypic changes than small variants [4-7]. The importance of SVs has been highlighted by their contribution to human diseases including cardiovascular diseases [8], autism [9], and a range of disorders [10]. Therefore, it is crucial to systematically profile SVs in the human genome for both biological research and clinical studies.

There are no gold-standard benchmarking procedures for SVs from next-generation sequencing (NGS) platforms. SVs from NGS are largely inferred from indirect evidence of disturbance of read mapping around the variation. Because SVs tend to reside within repetitive DNA and often span more base pairs than short reads (<1000 bp), the short reads of NGS usually lack sensitivity, leading to inevitable challenges in SV detection [11, 12]. Moreover, SV detection approaches vary in both sensitivity and specificity, as they emphasize on different SV-dependent and library-dependent features. Accurate identification of structural variation is very complex that requires characterization of the multifaceted features of SVs, including sequence information, type of variation, length, and location of breakpoints. As a result, different SV callers make inconsistent predictions [12, 13]. Therefore, owing to the complexity of SVs and the inconsistency of different SV callers, a comprehensive assessment of SV detection has been problematic.

Several efforts have been made in the community to benchmark SV calls. The Genome in a Bottle Consortium (GIAB) started building high-quality benchmark SV calls by distributing a set of 2,676 high-confidence deletions and 68 high-confidence insertions using SVClassify for a pilot genome (HG001/NA12878) in 2016 [14]. Recently, GIAB released a more comprehensive SV benchmark set for the son (HG002/NA24385) in an Ashkenazi Jewish trio with 2.66 Gb benchmark regions and 9,641 high-confidence SVs supported by at least one diploid assembly, but the identified SVs were not validated using experimental methods such as Sanger sequencing [15]. A well-characterized SV benchmark set is valuable in identifying false positive and false negative SVs called by various platforms and approaches. Yet, so far we don’t have an Asian-specific SV benchmark. The gnomAD-SV that comprises SVs from 14,891 genomes, revealed that different continental populations exhibited different levels of genetic diversity and SV features [16]. Therefore, designing an Asian benchmark is very necessary for promoting Asian genomic and disease research.

Our work is aimed at designing an Asian reference material comprising identified benchmark regions and high-confidence SV calls. This Asian benchmark is valuable for Asian studies in three aspects. Firstly, it provides material basis for Asian genomic and clinical research. It collects and preserves Asian genetic resources, accessible for Asian-specific biological experiments and drug screening. Secondly, the high-confidence SV calls for a benchmark cell line will serve as a gold-standard for evaluating the performance of a variety of SV detection platforms or strategies, including NGS and long-read sequencing technologies. Thirdly, this set of standards will become a threshold for clinical testing. It helps validate SV detection approaches in clinical practice, and on basis of the design of this benchmark, future benchmark comprising pathogenic SVs will be developed for clinical evaluation of SV detection related to specific diseases.

Establishment of immortalized cell lines is a routine strategy for building a reference material for biological research and clinical practice, among which immortalized B lymphocyte line transformed by Epstein-Barr virus (EBV) is a mainstream approach used by the international genetic storage institutions, including the NIGMS Human Genetic Cell Repository and the UK Biobank. EBV infection leads to B lymphocytes proliferation and immortalization *in vitro*, resulting in establishment of the immortalized B lymphocytes cell lines, which potentially provide unlimited genomic DNA resources and have been extensively used as a biological source for genetic and medical studies [17]. Previous studies and publications suggest EBV exists in the episomal form and is not integrated into the host cell chromosome, maintaining the host genome intact [18-20].

The advent of long-read sequencing technologies has greatly aided SV characterization. Although different long-read sequencing platforms applied diverse technologies, they are different from NGS by producing very long reads (1-100kb). In contract to NGS short-read data, the long-read data provide an advantage potential to increase the reliability and resolution of SV detection [21]. Given the advantages of long reads, our work established a high-confidence Asian SV benchmark for deletions and insertions by establishing an EBV immortalized B lymphocyte line and undertaking large-scale SV benchmarking across a range of the latest long-read sequencing or optical mapping techniques, including PacBio continuous long reads (CLR), PacBio circular consensus sequencing (CCS) reads, Oxford Nanopore long reads, and Bionano optical mapping (**Figure 1**). After comparing the performance of different platforms, we integrated and genotyped the final SV callset. Sanger sequencing validated the high confidence of our SV calls. We assembled haplotype-resolved diploid genomes by a trio binning approach using the PacBio CCS reads, and only high-confidence SVs supported by at least one diploid haplotype assembly will be included in the SV benchmark. The established cell lines and SV benchmark will provide a standard for assessing the precision and accuracy of different SV detection approaches, and ensure delivering accurate and reliable results for biological and genomic research in Asians. The immortalized B lymphocyte line will serve as an unlimited resource of Asian genomic DNA that can be extensively used as a biological source for future SV and medical studies.

**Figure 1.**
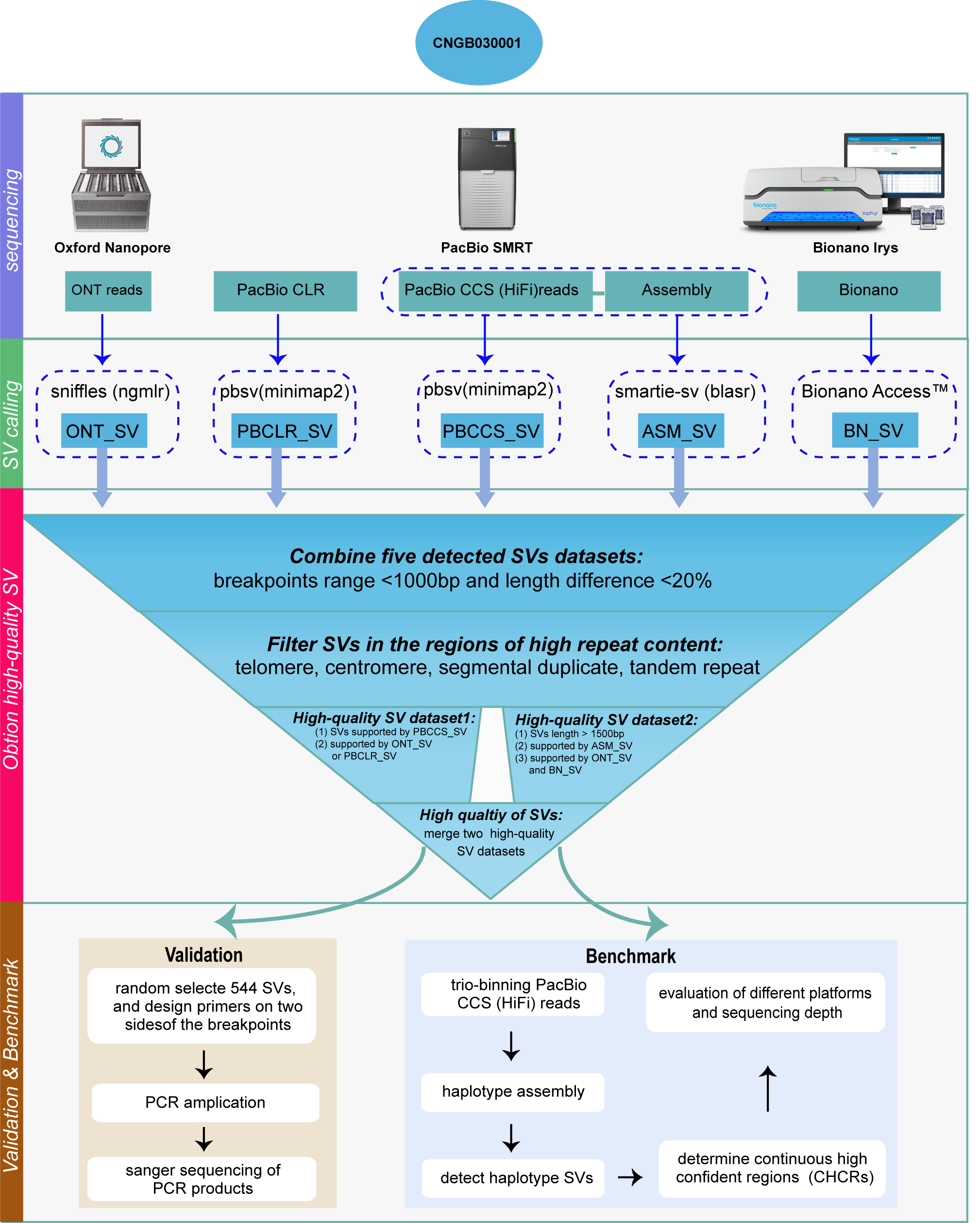
Workflow of establishing the benchmark SV set integrating different long-read sequencing technologies and approaches

## Results

### Sample establishment and sequencing results

Infection with Epstein-Barr virus (EBV) led to B lymphocytes proliferation and subsequent immortalization *in vitro*. Specifically, peripheral venous blood B lymphocytes of a normal Chinese man from Beijing were infected with EBV and treated with cyclosporine A (CyA) to increase the immortalization efficiency [17]. Morphology of the transformed cells was checked, and transformed cells were passaged and frozen for storage. Resuscitation experiments showed typical cell deformation and clonal growth characteristics. Free from contamination, cells were passaged continuously and grew well after resuscitation. Finally, an immortalized B lymphocyte line (CNGB030001) was successfully established.

By sequencing the cells, we generated 312.77Gb (∼104-fold) ONT data, 326.98 Gb (∼109-fold) PacBio CLR data, and 341.67 Gb (∼114-fold) Bionano data (**Table 1**). Compared with PacBio CLR, ONT showed a similar sequencing accuracy rate and obviously longer read length (CLR: 9.2 kb versus ONT: 24.6 kb). In addition, we obtained ∼22× highly accurate and long PacBio HiFi CCS reads after error correction from 869.48 Gb raw data (∼290×). The percentage of Q20 (accuracy rate: 99%) of the total CCS reads was 67.6% with an average read length of 12.0 kb, providing a high-quality foundation for SV calling. According to read length-GC plots, these four platforms performed very well in terms of uniformity in read length and GC content (Additional file 1: Figures S1-S4).

**Table 1.** Summary of sequencing results for different platforms.

| **Platform** | **Cell num** | **Sequencing type** | **Total bases (Gbp)** | **Depth (X)** | **Read length (avg. ± s.d., bp)** | **Read accuracy** | **GC content (%)** |
| --- | --- | --- | --- | --- | --- | --- | --- |
| PacBio CLR | 31 | subreads | 326.98 | 109 | $9,212 \pm 6,984$ | 0.8<sup>†</sup> | $42.66 \pm 5.82$ |
| PacBio CCS | 24 | CCS | 869.48; 71.85* | 266; 22X* | $11,961 \pm 3,662$ | $0.987 \pm 0.019$ | $40.59 \pm 5.63$ |
| Oxford Nanopore | 8 | 1D2 | 312.77 | 104X | $24,589 \pm 22,597$ | $0.849 \pm 0.028$ | $40.79 \pm 5.55$ |
| BioNano | 1 | BspQ1 | 341.67 | 114X | - | - | - |

*Note:* \* Total bases and depth after error correction for Circular Consensus Sequence (CCS). <sup>†</sup> Base quality of Pacbio CLR subreads produced by newer PacBio Sequel Sequencer is not available, so that we use an empirically overall read quality of 0.8.

### Candidate SV calls from different sequencing platforms

High accuracy is the prerequisite for establishing SV benchmark. For accuracy concern, we focused on detecting and characterizing large insertions and deletions in this work (Figure 1). By aligning PacBio CLR subreads, we identified 5,871 deletions and 6,936 insertions (**Table 2**). The size distribution of deletions displayed 300 bp and 6kb peaks related to SINE-Alu and LINE elements respectively (Additional file 1: Figure S5), suggesting effective SV calling by long reads [22, 23]. Compared to PacBio CLR, 17,901 SVs with 8,317 deletions and 9,584 insertions were obtained by aligning PacBio CCS sequencing reads (Table 2, Additional file 1: Figure S6). Most of the additional SVs from PacBio CCS were 50∼100 bp deletions. Similar to PacBio CLR result, both SINE-Alu and LINE deletions were identified, but no LINE elements for insertions were found in CCS SV calls, probably due to the limitation of read length (Additional file 1: Figure S6).

**Table 2.**
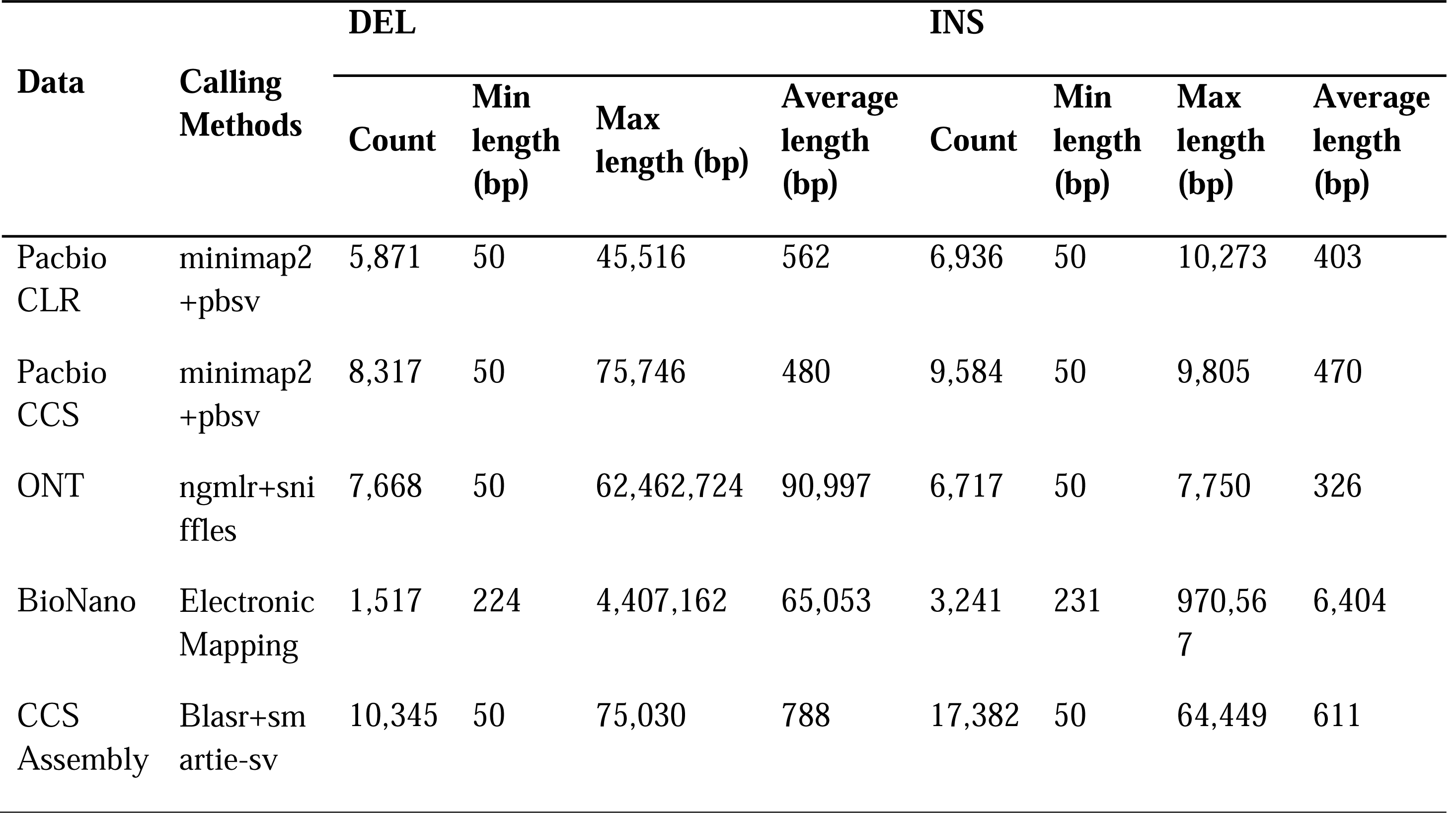
Insertion and deletion variation for different calling approaches.

The average read length of ONT data is longer than that of PacBio and the Bionano optical mapping relies on the density of restriction sites on the genome [24], thus theoretically they could efficiently detect the 6kb LINE elements for insertions. We detected 14,385 SVs with 7,668 deletions and 6,717 insertions using ONT data, as well as 4,758 SVs by Bionano (Table 2). A ∼6kb LINE insertion peak was successfully detected by both ONT and Bionano (Additional file 1: Figure S7-S8), but Bionano failed to detect the two short SINE-Alu events for deletions and insertions.

Apart from the alignment strategies, a *de novo* assembly-based method was applied for SV calling. We performed *de novo* assembly using 22× PacBio CCS reads, producing 3,542 contigs with the maximum length of 72 Mb and the N50 of 13 Mb. A good collinearity was observed from aligning the assembled contigs against the hs37d5 reference genome, indicating no visible structural errors were introduced in the assembly (Additional file 1: Figure S9). Finally, we detected 27,727 SVs using smartie-sv [25], more than those from alignment-based approaches (Table 2). The increase was mainly from small-scale insertions and deletions. Most noteworthy, the expected four insertion and deletion peaks related to SINE-Alu and LINE elements were all observed in the assembly-based SV calls (Additional file 1: Figure S10).

### Comparisons among different platforms

None of the approaches is comprehensive in discovering SVs on their own. A significant fraction of the identified variants was unique to a particular approach. The counts of unique SVs and SV overlapping among SV calls from different calling approaches were summarized in **Figure 2a**. PacBio CLR possessed the least unique variants (491), and CCS assembly-based SV calls had the most unique variants (7,930). Due to the specificity of Bionano, which is not accurate at base resolution, there were only 160 common SVs shared by all five calling approaches. With Bionano excluded, the three alignment-based single-molecular sequencing (PacBio CLR, PacBio CCS reads and ONT) approaches showed high consistency of SV calls with 8,156 common SVs. After integrating the CCS assembly-based result, the total number of common SVs reached 6,355 for the four datasets.

**Figure 2.**
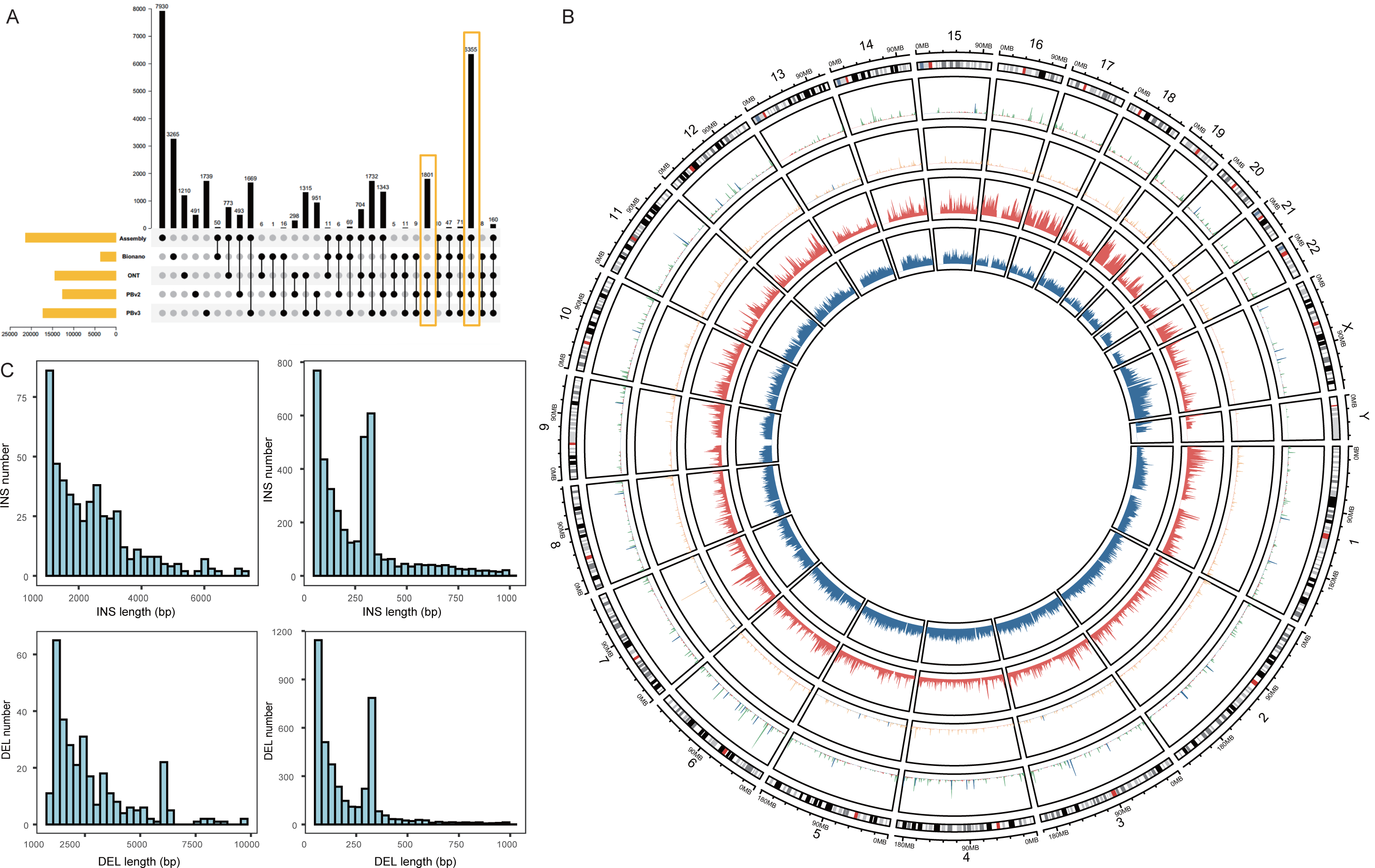
Comparisons among candidate SV call sets from different technologies and characterization of the high-confidence SV set. **A.** The number of SVs overlapping between candidate call sets from multiple approaches. **B.** A circos plot representing SV and repeat number in high-confidence SV calls using sliding non-overlapping windows of 1Mb length through the human genome. From the outer circle to the inner circle, the four circles represent the number of DEL, INS, SINE/Alu, LINE per 1 Mb window, respectively. **c** Size distribution for insertions and deletions in the high-confidence SV set.

### Construction of high-confidence SV calls

We integrated above candidate SV calls to construct a high-confidence SV callset by following specific steps and criteria (Figure 1). In consideration of the features of different sequencing platforms, there were two main reasons for applying these criteria. First, the outstanding long-read sequencing capacities of ONT, PacBio CLR, and Bionano guaranteed the longest possible read length, facilitating successful cover of large SVs. Second, PacBio CCS reads and CCS assembly approach emphasized on the high accuracy of the SVs. The longest possible read length and high accuracy guaranteed the high confidence of final SV calls. After filtering and integrating, a callset comprising 8,938 high-confidence SVs was established. SV distributions across all autosome chromosomes showed that the number of the distributed SVs had a good linear correlation with the length of the corresponding chromosomes (R^2^=0.85, p-value<0.0001, Additional file 1: Figure S11).

We examined the support for high-confidence SV callset from different sequencing platforms. Bionano showed the lowest support with 250 shared SVs compared to the high-confidence SV calls, and alignment-based CCS displayed the highest support with 8,914 shared SVs. The CCS assembly-based approach (6,419), PacBio CLR (6,797), and ONT (7,603) showed similarly high support (Additional file 1: Figure S12). The length distributions of high-confidence SVs clearly revealed four SINE-Alu and LINE peaks (Figure 2c). Moreover, distributions of the insertions and deletions on each chromosome were consistent with the density of SINE-Alu and LINE elements (Additional file 1: Figure 2b, Table S1).

### PCR and Sanger sequencing validated high-confidence SVs

To validate the accuracy of high-confidence SV calls, 400 SVs were randomly selected from 8,938 SVs for PCR amplification, of which 244 SVs were successfully amplified. We next randomly selected a second batch of 200 SVs that contained 56 amplification-failed SVs from the first batch and 144 new SVs. This time 22 of the 56 amplification-failed SVs were successfully amplified after PCR primer re-designs. Of the 544 SVs assessed by PCR, 203 and 341 are located in the genic and inter-gene regions, respectively. In total, 360 SVs were successfully amplified, and the overall amplification success rate was 66.2%, with amplification rate in genes reaching 90.6% – notably higher than 51.6% in inter-gene regions (**Figure 3B**). This result is not unexpected, as PCR amplification tends to be hindered by complex regions, such as repetitive abundant regions. Moreover, we analysed the length of amplified SVs and found smaller-size SVs had higher amplification rates than that of larger-size SVs (Additional file 1: Figure S13).

**Figure 3.**
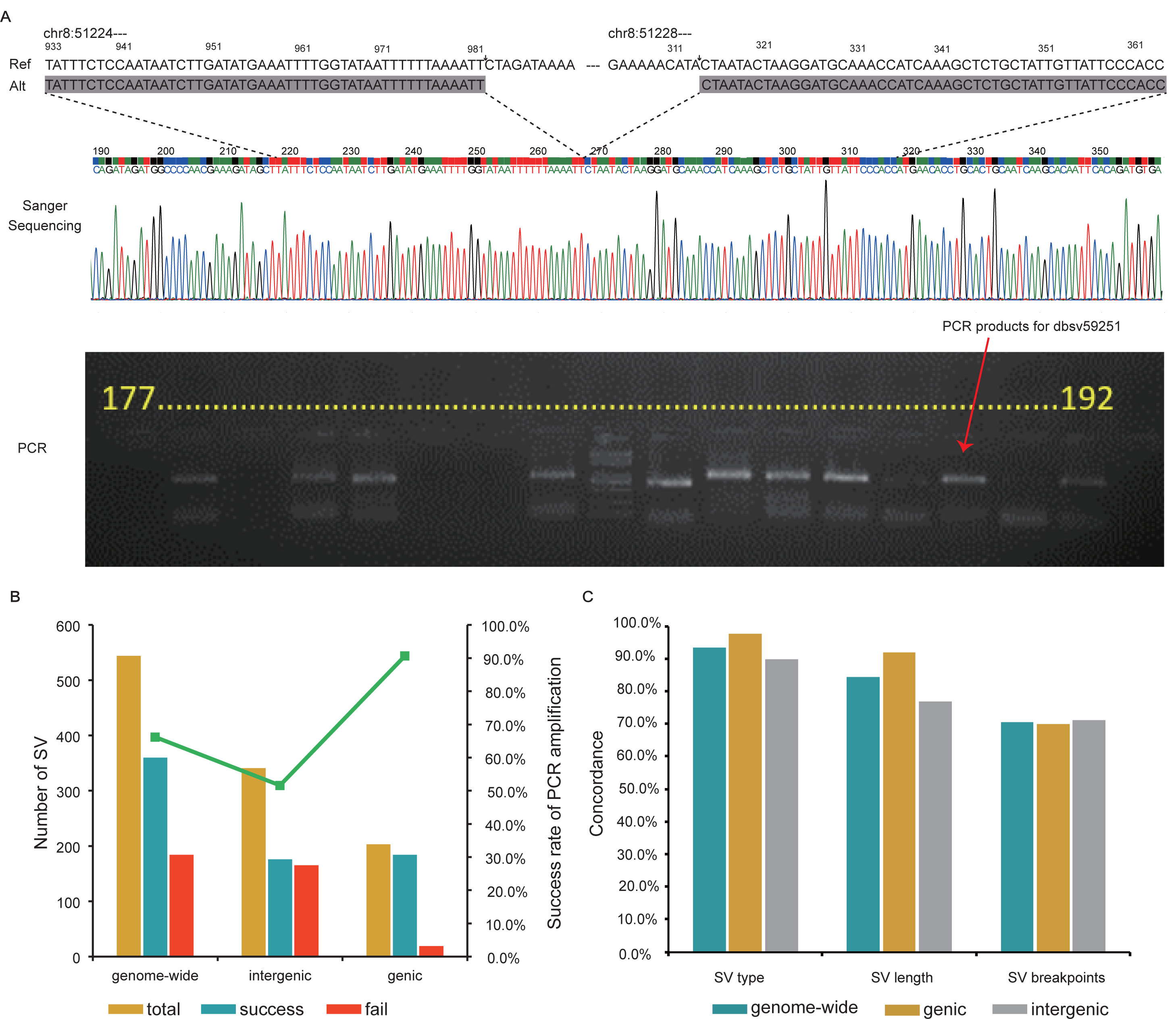
PCR and Sanger sequencing validated the SVs. **A.** An example of Sanger sequencing validating a deletion event on chromosome 8. **B.** PCR amplification rates for different genomic regions. **C.** Consistency rates of SV type, SV length, and SV position between Sanger sequenced variants and identified structural variants.

Among 360 amplification sites, 317 (∼88.1%) were successfully sequenced by paired-end Sanger sequencing and aligned to the reference genome (Figure 3A). The sequenced SVs were compared to the high-confidence SVs for evaluation by checking the SV type, SV length, and breakpoint position separately (Additional file 1: Figure S14). The consistency of length or breakpoint position was assessed by stringent matching of coordinate positions within 10 bp. Loci due to heterozygosity or low sequencing quality could not be effectively distinguished and were classified as uncertain. For instance, insertions exceeding 500 bp could not be detected by a single Sanger reaction, and thus were classified as uncertain. After excluding the uncertain sites, the concordance of SV type, length, and breakpoint position between Sanger sequenced SVs and our high-confidence SVs reached 93.5%, 84.3%, and 70.5%, respectively (Figure 3C). SVs in genic regions displayed higher concordance of type (97.7%) and length (91.9%) than SVs in intergenic regions (89.9% for type; 76.8% for length; Figure 3C). While the concordances of SV type and breakpoint position were not influenced by the SV size, the concordance of SV length dropped a little bit as SV size increased (Additional file 1: Figure S15). These results suggest that the high concordance from Sanger sequencing highly supports the robustness of our high-confidence SV calls.

### Establishment of the SV benchmark

With 8,938 high-confidence SV calls, we were aimed to construct a benchmark SV callset that could confidently exclude false positives and false negatives of the tested technology in the benchmark regions. To realize that, we applied a trio-binning based approach using the PacBio CCS data to identify haplotype-resolved SVs via haplotype assemblies, acting as another standard for proofing benchmark SVs. Specifically, using 183.6 Gb (61-fold) short-reads of the subject’s father and 184.7 Gb (62-fold) short-reads of the subject’s mother generated by DNBSEQ-G400 sequencing platform, 77.46% of the subject’s PacBio CCS data were unambiguously partitioned into paternal- and maternal-inherited reads using the trio-binning strategy by integrating five different *k*-mers [26] (Additional file 1: Table S2). Then we assembled two haplotypes using the biparental CCS reads by Canu independently [27]. The paternal and maternal haplotype assemblies spanned 2.76 Gb (contig N50 of 726 kb) and 2.92 Gb (contig N50 of 1,489 kb), respectively. The haplotype assemblies were aligned against human reference genome using blasr (v5.3.3) and SVs were called by smartie-sv independently.

On basis of haplotype assemblies, we constructed the continuous high-confidence regions (CHCRs), on which identified SVs should be arbitrarily supported by both the high-confidence SV calls and the paternal or maternal, haplotype-resolved SVs (**Figure 4A**). Finally, we identified 4,388 such CHCRs spanning 1.46 Gb with 6,882 high-confidence SV calls. These 6,882 SV calls constituted our final benchmark SV callset, serving as a gold-standard containing comprehensive SVs in benchmark genomic regions in Reference Material CNGB030001. In other words, in these 4,388 benchmark regions, we consider only those 6,882 benchmark SVs are expected in sample CNGB030001, which can be used to assess the performance of SV detection from different platforms and approaches.

**Figure 4.**
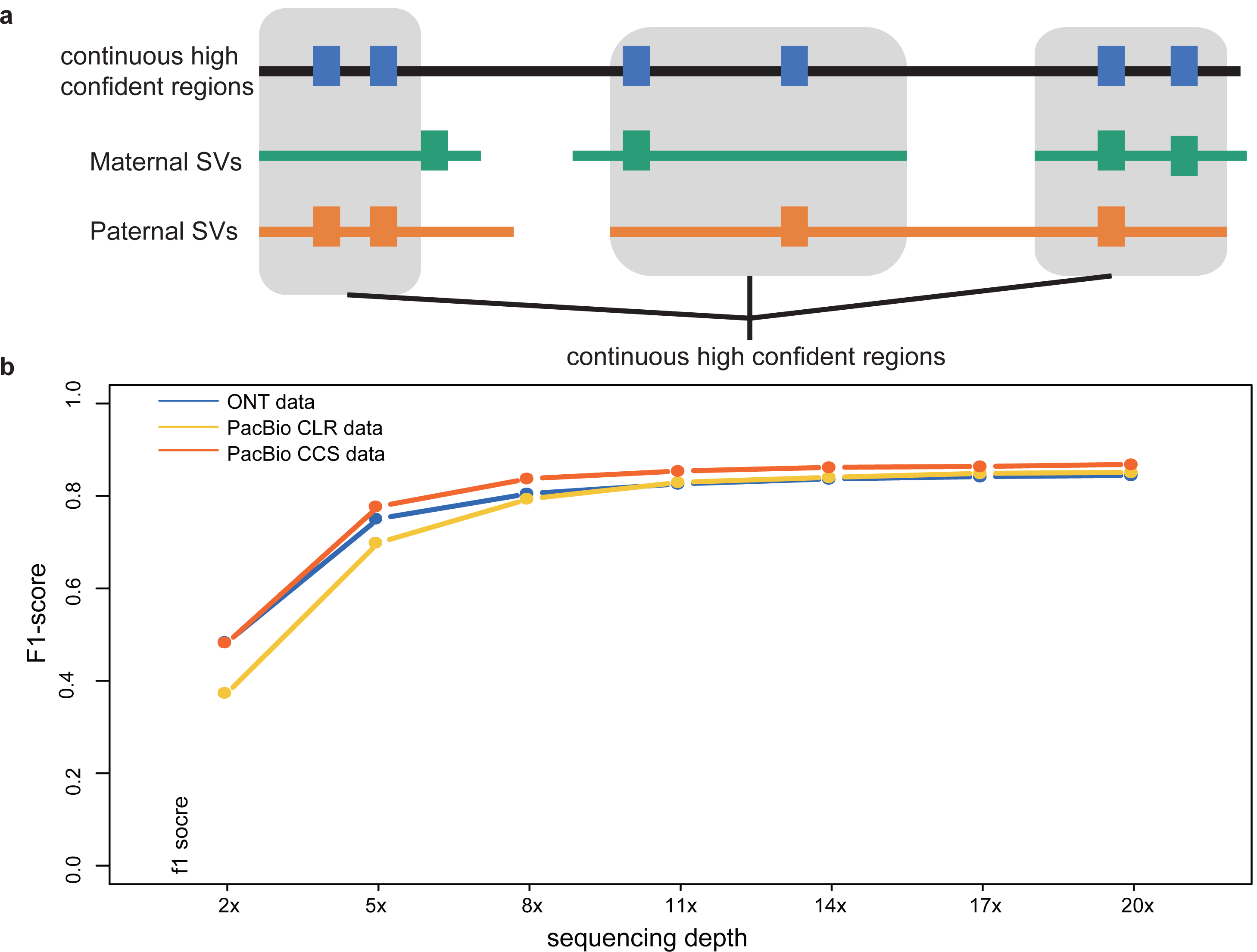
Assess the SV calling performance of different technologies. **A.** Construction of continuous confident regions based on two haplotype-resolved diploid SV callsets. **B.** Evaluation of different sequencing techniques and sequencing depth using the SV callsets.

### Comparison to GIAB benchmark

We compared our Asian benchmark to the recently released GIAB Tier 1 benchmark. Both benchmarks are designed for evaluating the detection of deletions and insertions. Our Asian benchmark comprises 3,346 deletions and 3,536 insertions, in contrast to the GIAB benchmark that includes 4,069 deletions and 5,262 insertions. In total, 3,326 SVs (48.3%) were shared by these two benchmarks. Comparison of benchmark SVs is dependent on the identified benchmark genome regions. When comparing the benchmark regions, 1.33 Gb of the 2.51 Gb benchmark regions in GIAB overlapped with our Asian benchmark regions. Within these overlapping regions, 3,313 SVs (62.4%) were shared by those two benchmarks, and 1,997 (37.6%) and 1,785 (35.0%) unique SVs were possessed by our Asian benchmark and GIAB benchmark, respectively (Additional file 1: Figure S16). This high overlap supports previous observations that many SVs are shared between different individuals [28]. Compared to common SVs, more small-size deletions and more SINE-Alu insertions were identified in Asian-specific SVs (Additional file 1: Figure S17). The unique SVs in overlapping benchmark regions reflect the genetic diversity of different individuals from different continents, which further illustrates the necessity of establishing Asian-specific reference materials and benchmarks.

### Application of the benchmark

The SV benchmark enables evaluation of the performance of different technologies in SV detection. Here we used our 6,882 benchmark SVs to assess the robustness of the three long-read sequencing technologies (PacBio CCS, CLR, and ONT) by checking their F1-scores under different sequencing depths. F1-scores of all sequencing platforms increased as their sequencing depth increased. When the sequencing depth reached to 11×, all F1-scores approached their saturation points (CCS: 85.4%, CLR: 83.0% and ONT: 82.6%) (Figure 4B). It should be noted that at higher sequencing depth (20×), PacBio CCS data was the top performer with a higher F1-score of 86.8% than the other two technologies (CLR: 85.1% and ONT: 84.5%). Our SV benchmark was constructed by integrating SV calls from diverse long fragment sequencing platforms and SVs from diploid assemblies. The F1-score results indicate that none of the three platforms could perfectly detect all SVs in benchmark regions on their own. It is necessary to integrate various approaches and technologies to realize comprehensive and confident SV detection.

We compared our benchmark SVs to the insertions and deletions identified by NGS data, which was generated in another parallel project on the same cell line CNGB030001, to evaluate the performance of NGS platforms in SV detection. We evaluated four representative detection tools including Manta [29], GRIDSS [30], LUMPY [31], and BreakDancer [32] to call SVs from MGISEQ-2000 and NovaSeq 6000 sequencing data, separately. Of the 3,536 insertions in our Asian benchmark, (a) 39.5% and 6.7% were detected by Manta and Gridss in MGISEQ data, respectively; (b) 28.3% and 14.3% were detected by Manta and Gridss in NovaSeq data, respectively; and (c) Lumpy and Breakdancer were not capable of detecting insertions from NGS data (Additional file 1: Table S3). Of the 3,346 deletions, (a) detection by Manta from MGISEQ and NovaSeq had 56.8% and 62.8% sensitivity, respectively; (b) Gridss, Lumpy, and Breakdancer displayed similar sensitivity in MGISEQ ranging from 38.3%∼ 41.4% and in NovaSeq ranging from 30.9%∼50.5% (Additional file 1: Table S3). These results suggest that SVs from NGS platforms differ among detection tools by displaying different sensitivities and specificities, and in general show low sensitivity to the detection of insertions. SV detection capacity also differs by different NGS platforms (MGISEQ-2000 *vs.* NovaSeq 6000). These results agree with the fact that different detection strategies emphasize on different SV-dependent and library-dependent features, highlighting the need for establishing SV benchmark using long reads.

## Discussion

A robust SV benchmark in specified genome regions provides gold-standard for evaluating the performance of various SV detection strategies and platforms in routine and clinical research. To form robust high-confidence SV calls, multiple SV call sets from a variety of methods and sequencing technologies need to be evaluated and integrated. PacBio adopts a sequencing-by-synthesis strategy and produces two types of reads. The continuous long reads (CLR) emphasizes on the longest possible reads, and the circular consensus sequencing (CCS) reads are featured for high accuracy (> 97%). Oxford Nanopore Technology (ONT) works by monitoring changes to an electrical current as nucleic acids are passed through a protein nanopore. The Bionano linearizes and images long DNA strands that are nicked and fluorescently labeled to produce single molecule physical maps.

Our work established 8,938 high-confidence SV calls by combining SV call sets using both alignment-based SV callers and *de novo* assembly-based SV callers from above state-of-art long-read sequencing technologies, and applied experiments and haplotype assemblies for validating and establishing a robust benchmark SV callset. Compared to previous SV work, this study collected high-confidence SV calls by incorporating a deeply sequenced new data type, PacBio CCS sequencing, conducted experimental validation, and applied the new assembly approach trio binning for diploid *de novo* assemblies (combining two parental whole genome sequencing data) to establish robust benchmark SV calls.

We established an Asian SV benchmark for identifying both false negatives and false positives in specified benchmark regions with a well-characterized set of haplotype-resolved SVs. The final benchmark SV callset comprising 6,882 SVs is highly robust. Firstly, it was established on basis of 8,938 high-confidence SVs, which were constructed by integrating comprehensive information from the state-of-art long-read sequencing technologies. Different sequencing platforms and analysis approaches (alignment-based and assembly-based) complemented each other and an integration of them was robust in detecting confident SVs. All SVs in the high-confidence set had support in reads from more than one technology. Secondly, we validated randomly selected SVs using PCR amplification and Sanger sequencing, confirming the high confidence of our SV calls. Lastly, we took use of trio binning-based haplotype assemblies to distinguish paternal or maternal SVs. Only haplotype-resolved, high-confidence SVs could be included in the benchmark SV calls. The established Asian benchmark spans 1.46Gb and covers 6,882 SVs supported by at least one diploid haplotype assembly, allowing the community to confidently evaluate the detection capacity for insertions and deletions in future practices.

It should be noted that high accuracy is a prerequisite for establishment of the benchmark. Therefore, our established benchmark only covered specific regions of the genome with confirmed accuracy, not the whole genome. In those regions we confirmed that benchmark SVs were highly confident. Our benchmark did not focus on complex SVs (e.g., inversions, duplications, and translocations) either. Importantly, we emphasize that this SV benchmark allows the community to confidently evaluate the performance of various platforms and approaches in detecting insertions and deletions. The deeply sequenced data in this study can be used in future work to extend our understanding of complex SVs. As mentioned above, in another parallel project [33] we generated about 4.16T clean data of the same cell line using seven sequencing strategies in different laboratories, including two BGI regular NGS platforms, three Illumina regular NGS platforms, single tube long fragment read (stLFR) sequencing, and 10X Genomics Chromium linked-read sequencing. Those large data will provide comprehensive variants information, serving as valuable genomic resources facilitating future genomic or medical research.

By analysing extensive SV calls generated from different platforms and calling tools, we found that different technologies had distinct strengths and weaknesses. PacBio CCS reads detected ∼5000 more SVs than PacBio CLR, but neither of them identified the 6 kb LINE insertions using the alignment-based strategy. CCS assembly-based approach successfully identified four SINE/Alu and LINE elements in insertions and deletions, and detected the largest SV callset with more small-size insertions and deletions. Bionano mapping is based on optical ultra-long single molecules of DNA that are fluorescently labelled at specific restriction sites [24]. Due to its dependency on the density of restriction sites, it failed to accurately detect small-size SVs with the least SVs identified. While other techniques detected more insertions than deletions, ONT was more sensitive to detect deletions than insertions. It could also effectively detect the 6 kb LINE element insertions.

The established reference material CNGB030001 serves as an unlimited resource of Asian genomic resources, facilitating future Asian SV and medical studies. EBV transformed cell lines are widely used internationally in routine and clinical research. Usually cell line genome is relatively stable under a certain number of passages, and after long-term passages genomic instability is a common problem in immortalized cell lines, such as tumor cell lines. In our project, genomic instability after long-term passages wouldn’t be a concern for our cell line applications. To release as reference material, we have generated a large quantity of tubes at one time to confirm usage for several years, ensuring low cell passages. Good cell bank management could effectively ensure low cell generations, and regular cell line identifications will help verify the cell line stability. Therefore, cell line CNGB030001 can be widely used for Asian genomic and medical research as a valuable reference material.

## Conclusion

Taking advantage of multiple long fragment sequencing platforms, our work established an Asian reference material and developed a robust SV benchmark. PCR amplification and Sanger sequencing were conducted to validate the high quality of our high-confidence SVs. Haplotype assemblies via trio binning were used for identifying haplotype-resolved SVs to construct the final robust benchmark. The performance of SV calling of different technologies across various sequencing depths provides valuable information for further SV studies. Finally, we report that the established benchmark cell line provides valuable Asian genomic resources in biological and medical research and the SV benchmark can serve as a gold standard for benchmarking SV detection approaches in clinical practice.

## Materials and methods

### Establishment of immortalized B lymphocyte line

B lymphocyte immortalization was performed according to the published protocol [34] with slight modifications. 4.5 mL whole blood was collected from a Chinese donor with sodium citrate anticoagulant blood collection tube (369714, Becton Dickinson, New Jersey, US). Peripheral blood mononuclear cells (PBMCs) were isolated from the whole blood by Ficoll density gradient centrifugation (p-05824, GE Healthcare, Illinois, US). The lymphocytes were simulated and transformed by treating cyclosporin A (No.12088, Cayman Chemical, Michigan, US) and EBV that was prepared from collecting the supernatant of B95-8 cells (ATCC CRL-1612). The transformed lymphocytes were monitored by microscope for the performance of transformation. After transformation, the immortalized lymphocytes were cultured on large scale and then divided into 1 × 10^6^ per tube for long-term storage.

### DNA extraction, library preparation, and sequencing

#### DNA extraction for PacBio and Oxford nanopore

5 × 10^6^ frozen cells were suspended in 1 × PBS buffer to reach a total volume of 2 ml, and 1 volume of ice-cold cell lysis buffer (1.28 M sucrose, 40 mM Tris hydrochloride, 20 mM MgCl_2_, 4% Triton X-100 [pH 7.5]) and 3 volumes of ice-cold distilled water were added. The mixture was incubated for 10 minutes on ice, and nuclear pellets were collected by centrifugation (6,000 rpm, 5 min, 4°C). The nuclei were completely resuspended in extraction buffer (0.8M guanidine hydrochloride, 30mM Tris, 30mM EDTA, 5% Tween-20, 0.5% Triton-100 [pH 8.0]) containing 1% sodium dodecyl sulphate (SDS) and proteinase K (2 mg/ml final concentration), and incubated at 56°C for 2 hours. gDNAs were extracted by phenol-chloroform-isoamyl alcohol (25:24:1 by volume) and chloroform-isoamyl alcohol (24:1 by volume), and precipitated with 0.7 volume of isopropyl alcohol at −20°C for 40 minutes. The DNA precipitates were washed in ice-cold 80% ethanol for twice, collected by centrifugation (12,000 rpm, 15 min, 4°C), dried under vacuum, and finally resuspended in 100 ul of EB (10 mM Tris hydrochloride [pH 8.0]). To acquire high-quality DNA, an additional step of purification was performed right after DNA extraction by 0.8 volume of magnet beads from AMPure XP kit (#A63882, Agencourt) according to the manufacturer’s instructions. Agilent 4200 Bioanalyzer (Agilent Technologies, Palo Alto, California) was used to detect the integrity of gDNA. 8 μg of gDNA was sheared using g-Tubes (Covaris) and concentrated with the AMPure PB magnetic beads.

#### Library construction and sequencing of PaBio CLR

We used a Pacific Biosciences SMRT bell template prep kit 1.0 to construct each SMRT bell library by following the manufacturer’s instructions. The constructed libraries were size-selected on a BluePippin™ system for molecules ≥20 Kb, followed by primer annealing and the binding of SMRT bell templates to polymerases with the DNA/Polymerase Binding Kit. Finally, sequencing was performed on the Pacific Bioscience Sequel platform (Annoroad Gene Technology Co., Ltd China) for 10 h and finally sequenced by CLR mode with the Sequel System (Pacific Biosciences, CA).

#### Library construction and sequencing of PacBio CCS

SMRTbell libraries were prepared using the ‘Express Template Prep Kit 1.0’ protocol (Pacific Biosciences, CA). 5 ug genomic DNA was sheared to ∼15 kb fragments by a g-TUBE (#520079, Covaris) centrifugation (2,000 Xg, 2 min, twice), was size-selected for 10 kb using the BluePippin system (Sage Science, MA) by marker (0.75% DF Marker S1 High-Pass 6-10kb vs3) for the 10-20 kb DNA target fragments. Quality control of the libraries was performed by Qubit fluorometer (Life Technologies, CA) and Bioanalyzer 2100 (Agilent, CA). The prepared library was loaded into SMRT cell 1M by Sequel Binding Kit 3.0 (Pacific Biosciences, CA) and finally sequenced by CCS mode with the Sequel System (Pacific Biosciences, CA).

#### Library construction and sequencing of Oxford nanopore

Genomic DNA libraries were prepared using the Ligation sequencing 1D kit (SQK-LSK109, Oxford Nanopore Tech., UK). End-repair and dA-tailing of DNA fragments were performed using the Ultra II End Prep module (#E7546, NEB) by following protocol recommendations. The dA tailed sample was tethered to 1D adapter by Quick Ligation Module (#E6056, NEB). Finally, the prepared DNA library was loaded into FLO-PRO002 flow cell and sequenced on PromethION (Oxford Nanopore Tech., UK).

#### DNA extraction and sequencing for Bionano

The isolation of high-molecular-weight gDNA from immortalized B lymphocyte line was performed with the Bionano Prep™ Cell Culture DNA Isolation Kit according to the standard protocol of Bionano Prep Cell Culture DNA Isolation Protocol (Document Number: 30026). Sequence-specific labelling of megabase gDNA for Bionano mapping was conducted by nicking, labelling, repairing, and staining (NLRS) following the standard protocol of Bionano Prep™ Labelling-NLRS. The labelled gDNA was transferred into Bionano Genomics Saphyr™ for scanning to obtain the optical map.

### SV calling based on different platforms and methods

#### Alignment-based SV calling

(**a**) For CLR reads, BAM files of CLR reads were exported from SMRT Link (v6.0.0.47841), and aligned to the reference genome (hs37d5) using minimap2 (v2.15-r906-dirty) [35] with parameters “align_opts = “-x map-pb -a --eqx -L -O 5,56 -E 4,1 -B 5 --secondary=no - z 400,50 -r 2k -Y -R “@RG\tID:rg1a\tSM:human”“. (**b**) For CCS reads, CCS reads BAM files were aligned to the reference genome (hs37d5) using minimap2 (v2.15-r906-dirty) [35] with parameters “-R -t 2 --MD -Y -L -a -x map-pb”. According to the mapping positions, SAMtools (v0.1.19) [36] was used to sort the alignments with default parameters. To identify SV, pbsv (v2.1.1) [37] with default parameters was used to sort alignment files. (**c**) For ONT reads, reads with quality score > 7 were aligned to the reference genome (hs37d5) using ngmlr (v0.2.7) [38] with the parameter “--presets nanopore”. SVs were called using sniffles (version 1.0.8) [38] with parameters “--min_support 1 --threads 8 --num_reads_report -1 – genotype”. (**d**) For Bionano data, Bionano data was generated from enzyme BspQI and SVs were called using Bionano Solve pipeline (v3.1) [39] with default parameters.

#### De novo assembly-based SV calling

Falcon (v0.3.0) [40] was used for assembly and contigs were aligned to hs37d5 reference genome using blasr (v5.3.3) [41]. SV calling was performed with smartie-sv [25, 42].

### Integration of the high-confidence SV calls

The high-confidence SV callsets were integrated from all candidate SV callsets by the following processes: **(a)** The same type of SVs within 1kb with sequence change <20% were merged into a single SV using SVmerge (v1.2r27) [43]. (**b**) All SVs detected by PacBio CCS Reads and supported by either PacBio CLR or ONT were retained. SVs longer than 1.5 kb supported by both PacBio CCS assembly and Bionano mapping were also retained. **(c**) SVs located in centromeres, telomeres, segmental duplications, and short tandem repeat regions were removed according to the SV annotations by annovar (v20160201) [44]. (**d**) Hawkeye (v2.0) [45] was used for SV visualization by automatically outputting images for manual checking.

### SV validation by PCR amplification and Sanger sequencing

We performed the Sanger validation for two batches of randomly selected SVs. An amplification was considered successful if a clear single band was observed or the expected size band could be purified and separated by gel cutting. The failed SVs showed ambiguous bands. To evaluate the effect of primer design, failed SVs from the first batch were repeated for PCR amplification in the second batch. The corresponding primers were designed with Primer3 by default parameters [46]. Amplification results for each amplicon were validated by electrophoresis and the products were loaded onto 3730 sequencers with pair-end sequencing mode (Thermofisher, Massachusetts, US). Raw sequencing results were analysed by Sequence Scanner Software v2.0 and the low-quality parts were trimmed. The clean reads were mapped to the reference genome hs37d5 by BLAST, and the mapping results were manually checked for SVs. For manual curation, the following criteria were used to evaluate the accuracy of previous SV calls: 1) if there was an SV event supported by any single sanger read; 2) if a previously called SV could match a Sanger call within 10-bp difference in size; and 3) if the breakpoint of a previously called SV could match that of a Sanger call within 10 bp differences.

### Construction of diploid haplotype genomes using trio binning

Short reads from the parents were used to identify *k*-mers unique to each parent and partition (“trio binning”) the CCS reads. The trio binning pipeline was applied to partition paternal and maternal CCS reads [26, 47] using five different *k*-mers including 21 bp (previously reported for trio binning) and longer *k*-mers of 41, 51, 61 and 81 bp. To realize accurate partition, one integration method was used following two criteria: for one CCS read, (**a**) at least two different k-mers support the same parental source; (**b**) more than half of the different k-mers support the same parental source.

To obtain paternal and maternal haplotype genomes, we used canu (v1.8-r9528) [27] to assemble paternal and maternal CCS reads with parameters “-trim-assemble genomeSize=3100m correctedErrorRate=0.039 -pacbio-corrected”, separately. Meanwhile, unassigned CCS reads were used in both two assemblies. Lastly, the paternal and maternal haplotype assemblies were aligned against human reference genome using blasr (v5.3.3) and SVs were called by smartie-sv independently.

### SV benchmark construction, comparison, and application

To establish the benchmark SV callset, we identified the continuous high-confident regions by combining 8,938 high-confidence SVs and SVs called from haplotype-solved assemblies. To evaluate the capability of different platforms, SV calls detected by different technologies were all converted into VCF formats and evaluated against the continuous high-confident regions using Truvari (v1.3) [48] with default parameters. To compare our Asian benchmark to GIAB benchmark, we used SVmerge (v1.2r27) with default parameter “-d 1000” for SV positions, “-l 0.5” for SV length difference, and “-r 0.5” for SV overlap. Overlapping and unique SVs were enriched for gene pathways by R package clusterProfiler.

SVs from two NGS platforms (MGISEQ-2000 and NovaSeq 6000, triplicat calls per sample) were called using four tools and hs37d5 reference genome with corresponding parameters: GRIDSS (default parameters), LUMPY (default parameters), Manta (minCandidateSpanningCount=3, minScoredVariantSize=50, minDiploidVariantScore =10, minPassDiploidVariantScore = 20, minPassDiploidGTScore = 15, minSomaticScore =10, minPassSomaticScore=30, useOverlapPairEvidence = 0, enableRemoteReadRetrievalForInsertionsInGermlineCallingModes = 1, enableRemoteReadRetrievalForInsertionsInCancerCallingModes = 0), and BreakDancer (num:10001, lower:78.10, upper:465.35, mean:254.28, std:48.32, SWnormality:-31.28). For each sample, 3 SV calls were integrated into a final call by SURVIVOR using following parameters: 1000 2 1 1 0 30. We used SVmerge (v1.2r27) with default parameter “-d 1000” for SV positions, “-l 0.5” for SV length difference, and “-r 0.5” for SV overlap. To find common SVs between NGS SV results and our SV benchmark.

## Acknowledgments

We thank Inge Seim for reading and offering very helpful suggestions on the writing of this manuscript.

## Authors’ contributions

JH, DW, and FC conceived the project. JH, OW, GF, SQ, FL, and FC supervised the study. JH, LL, WZ, and JZ contributed to sample establishment. FL, SL, SS, YS, XL, WQ, WY, JZ, YX, and YW performed bioinformatics analyses. FF, OW, LW, FC, and SQ performed PCR amplification and Sanger sequencing for evaluation of structural variation. JH, XD, SS, GF, and OW wrote the manuscript with help from all co-authors.

## Availability of data and materials

The sequencing data (including ONT, PacBio CCS, and PacBio CLR data) has been deposited in the CNGB under accession number CNP0000091. Bionano data can be accessible at CNGB FTP site: http://ftp.cngb.org/pub/CNSA/data1/CNP0000091/.

## Ethics approval and consent to participate

The experimental procedures were in accordance with the guidelines approved by the institutional review board on bioethics and biosafety of BGI (IRB-BGI). The experiment was authorized by IRB-BGI (under NO. FT19038), and the review procedures in IRB-BGI meet good clinical practice (GCP) principles.

## Consent for publication

Not applicable.

## Supplementary material

**Additional file 1: Supplementary figures and tables.**

**Figure S1** The length and GC content distributions of sequencing reads produced from PacBio continuous long read (CLR) technology

**Figure S2** The quality of reads varies with the number of sequencing passes produced from PacBio circular consensus sequencing (CCS) technology

**Figure S3** The length and GC content distributions of sequencing reads produced from PacBio CCS technology

**Figure S4** The length and GC content distributions of sequencing reads produced from Oxford Nanopore technology (ONT)

**Figure S5** The size distributions for 0∼1000bp and 1000∼10000bp ranges for candidate SV calls from PacBio CLR

**Figure S6** The size distributions for 0∼1000bp and 1000∼10000bp ranges for candidate SV calls from PacBio CCS

**Figure S7** The size distributions for 0∼1000bp and 1000∼10000bp ranges for candidate SV calls from ONT

**Figure S8** The size distributions for 0∼1000bp and 1000∼10000bp ranges for candidate SV calls from Bionano

**Figure S9** Dot plot showed the synteny between assembled contigs from PacBio CCS reads and the hs37d5 reference genome

**Figure S10** Size distributions for 0∼1000bp and 1000∼10000bp ranges for candidate SV calls from PacBio CCS assembly

**Figure S11** The relationship between chromosome length and the number of SVs

**Figure S12** The number of unique SVs and SVs overlapping among different candidate SV callsets

**Figure S13** Distribution of PCR Amplification results in different length ranges

**Figure S14** Consistency rates of SV type, SV length, and breakpoint position between Sanger sequenced variants and high-confidence SV calls with uncertain sites not excluded

**Figure S15** Consistency rates of SV type, SV length, and SV position between Sanger sequenced variants and high-confidence SV calls in different length ranges. Uncertain sites were not excluded

**Figure S16** Comparison of the Asian CNGB030001 benchmark and the GIAB HG002 benchmark within the 1.33 Gb overlapping benchmark regions.

**Figure S17** The size distributions for 50∼1000bp and > 1kb ranges for common and Asian unique SVs by comparing GIAB benchmark to the Asian CNGB030001 benchmark.

**Figure S18** GO enrichment of Asian-benchmark-specific SVs.

**Table S1** Counts of identified high-confidence SVs on each chromosome

**Table S2** Trio binning of CCS reads

**Table S3** Counts of overlapping SVs from NGS platforms in comparison to the SV benchmark

