## Additional file 1 for "Robust Benchmark Structural Variant Calls of An Asian Using the State-of-Art Long Fragment Sequencing Technologies"

### Supplementary Figures

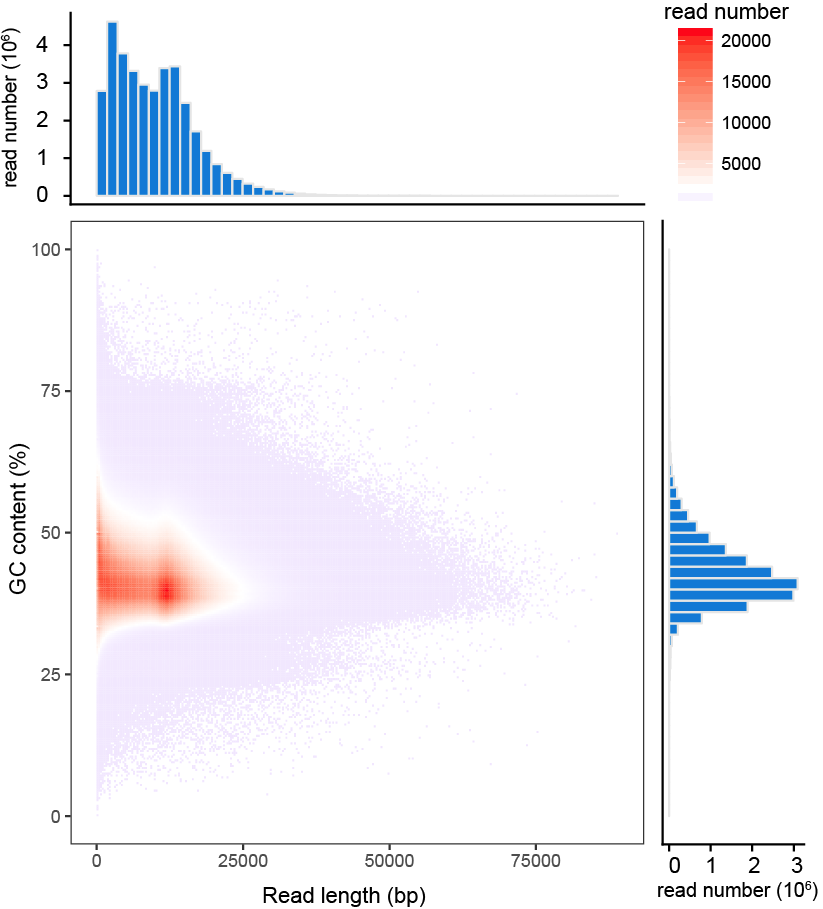

#### Figure S1 The length and GC content distributions of sequencing reads produced from PacBio continuous long read (CLR) technology

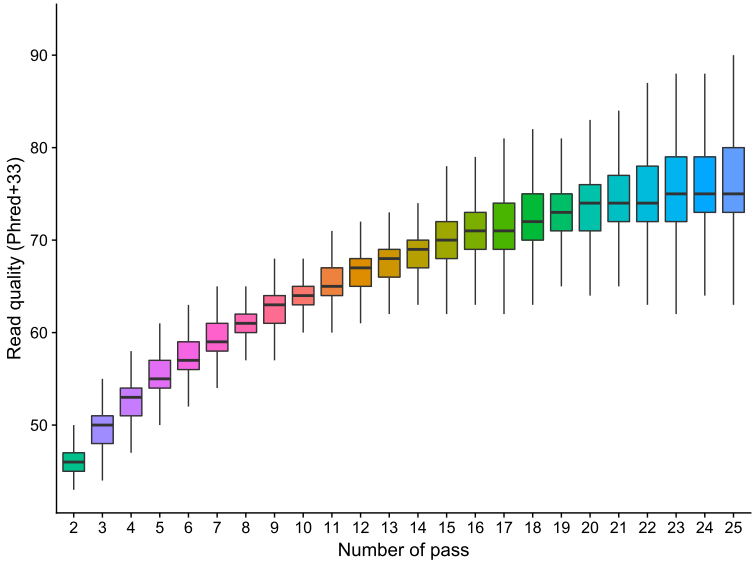

#### Figure S2 The quality of reads varies with the number of sequencing passes produced from PacBio circular consensus sequencing (CCS) technology

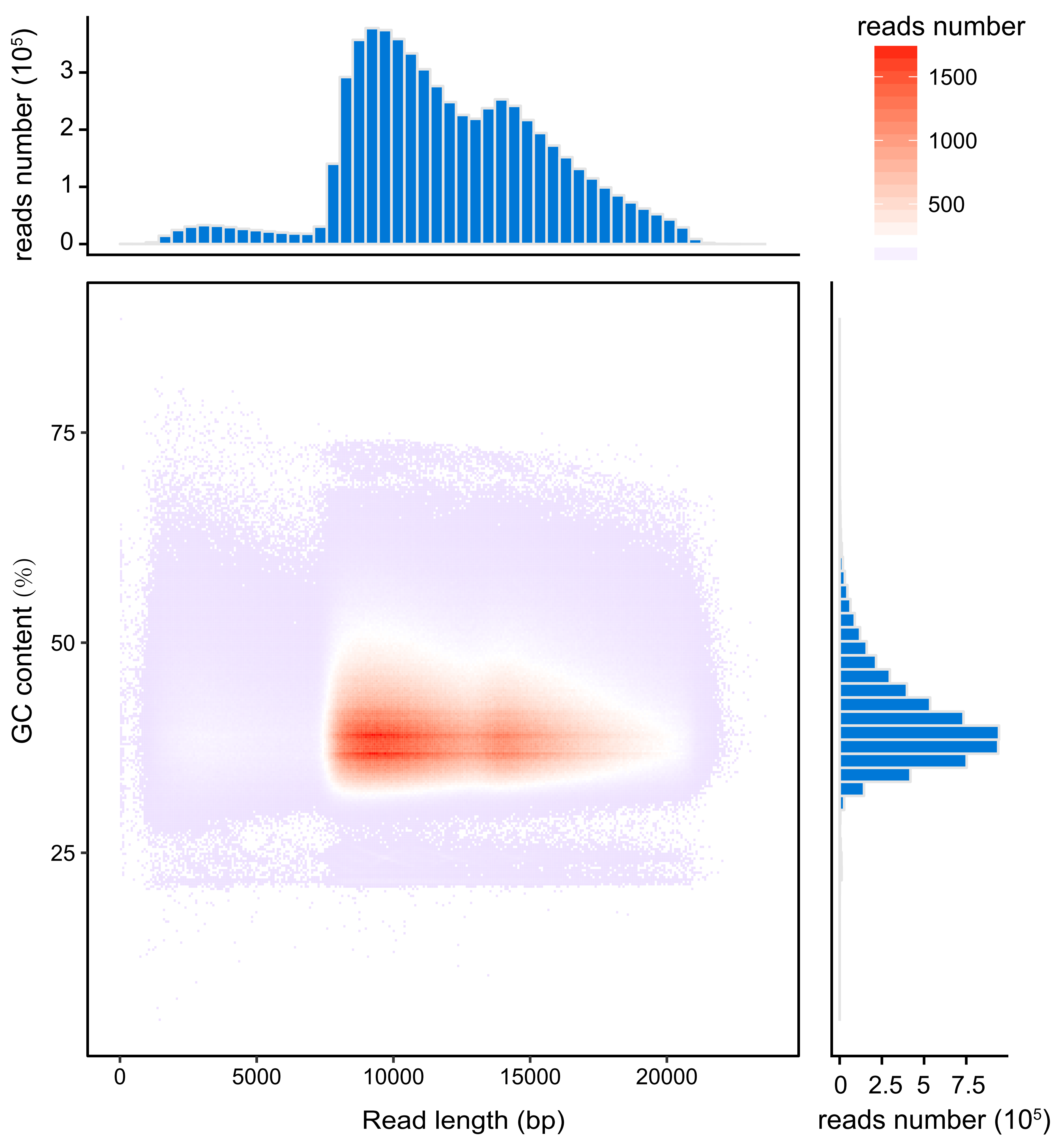

#### Figure S3 The length and GC content distributions of sequencing reads produced from PacBio CCS technology

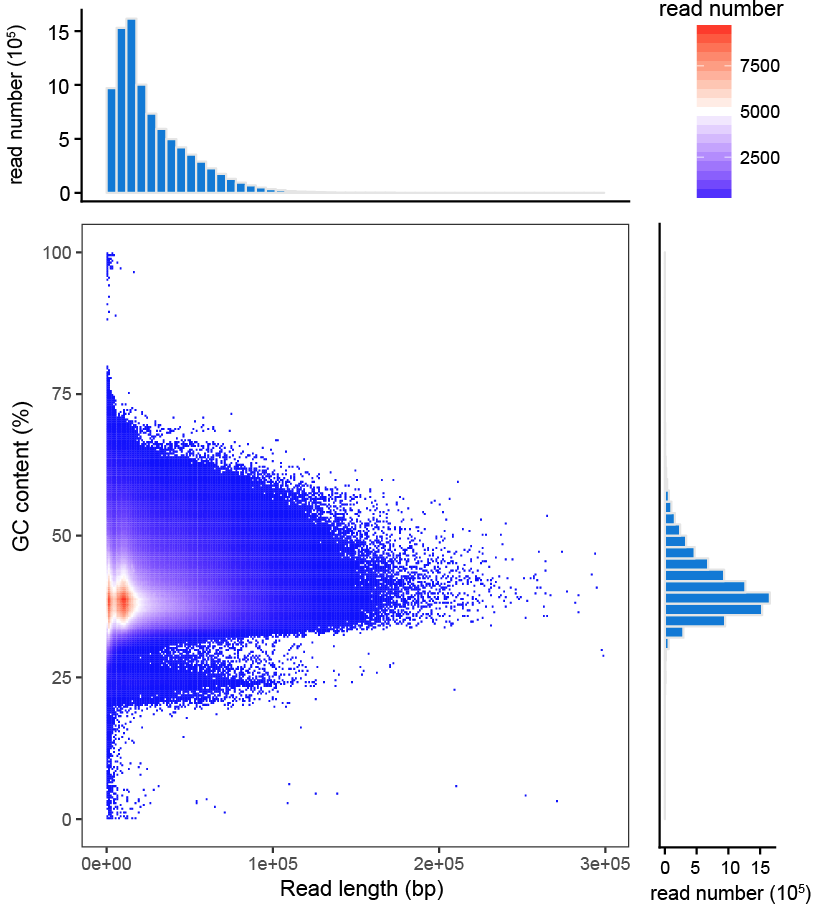

#### Figure S4 The length and GC content distributions of sequencing reads produced from Oxford Nanopore technology (ONT)

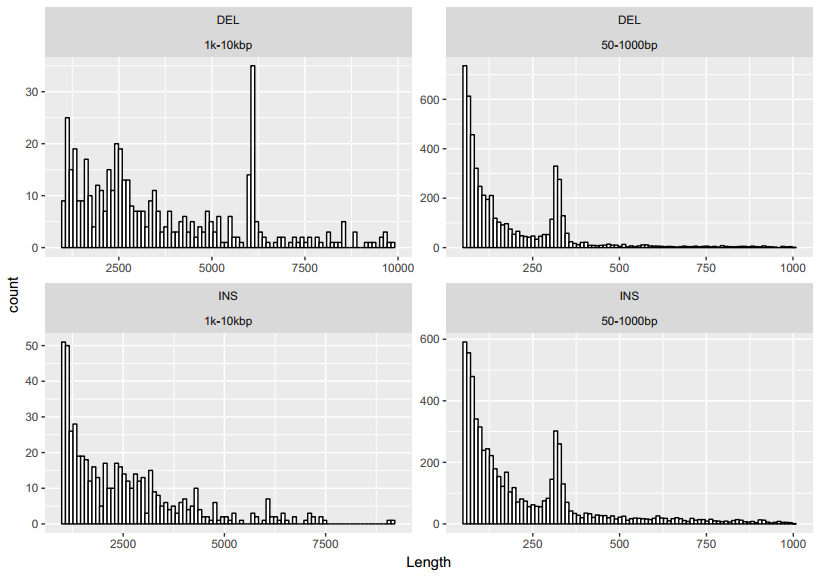

#### Figure S5 The size distributions for 0~1000bp and 1000~10000bp ranges for candidate SV calls from PacBio CLR. Alu and LINE insertion/deletion events are at around 300 bp and 6000 bp.

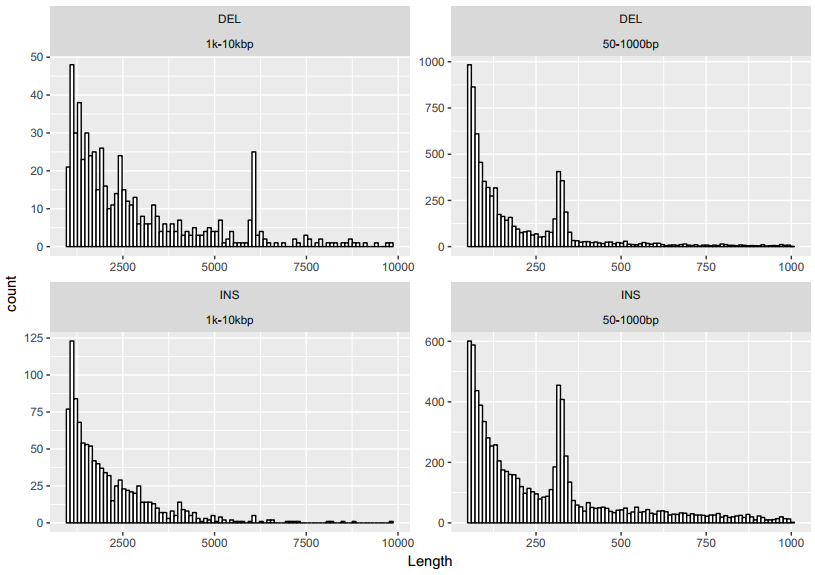

#### Figure S6 The size distributions for 0~1000bp and 1000~10000bp range for candidate SV calls from PacBio CCS. Alu and LINE insertion/deletion events are at around 300 bp and 6000 bp.

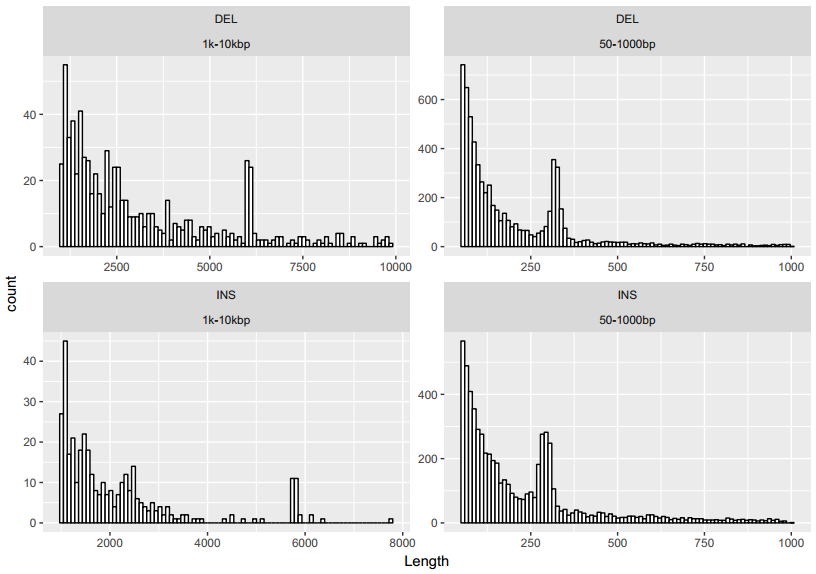

#### Figure S7 The size distributions for 0~1000bp and 1000~10000bp ranges for candidate SV calls from ONT. Alu and LINE insertion/deletion events are at around 300 bp and 6000 bp.

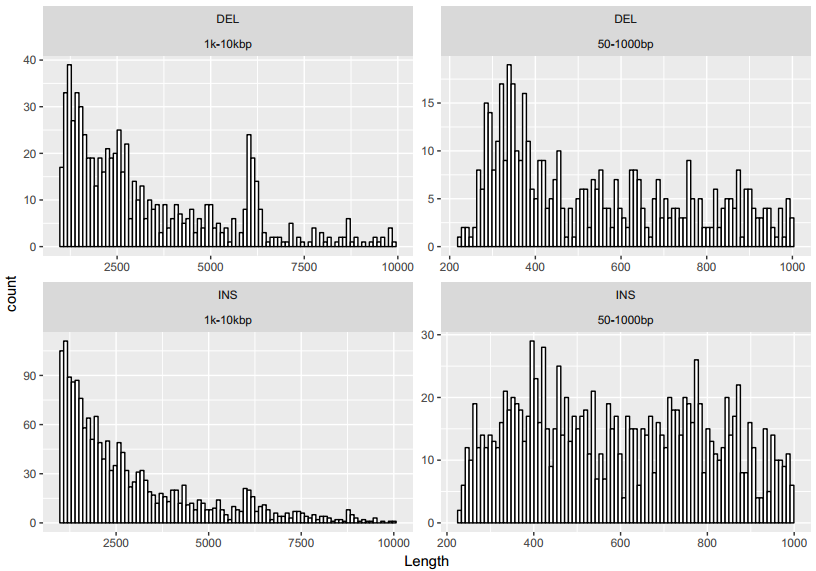

#### Figure S8. The size distributions for 0~1000bp and 1000~10000bp ranges for candidate SV calls from Bionano. Alu and LINE insertion/deletion events are at around 300 bp and 6000 bp.

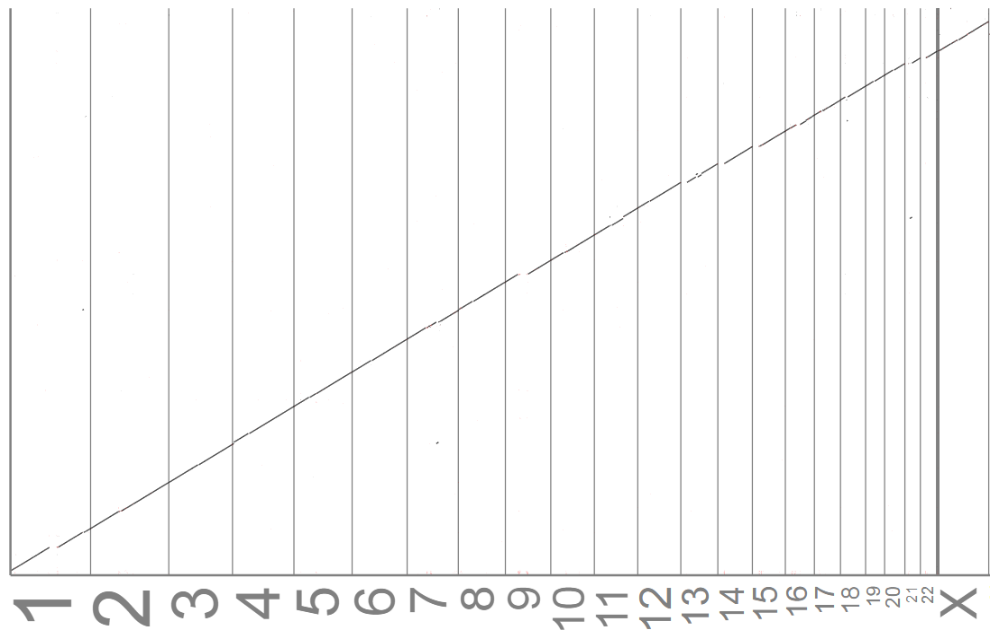

#### Figure S9 Dot plot showed the synteny between the assembled contigs from PacBio CCS reads and the hs37d5 reference genome

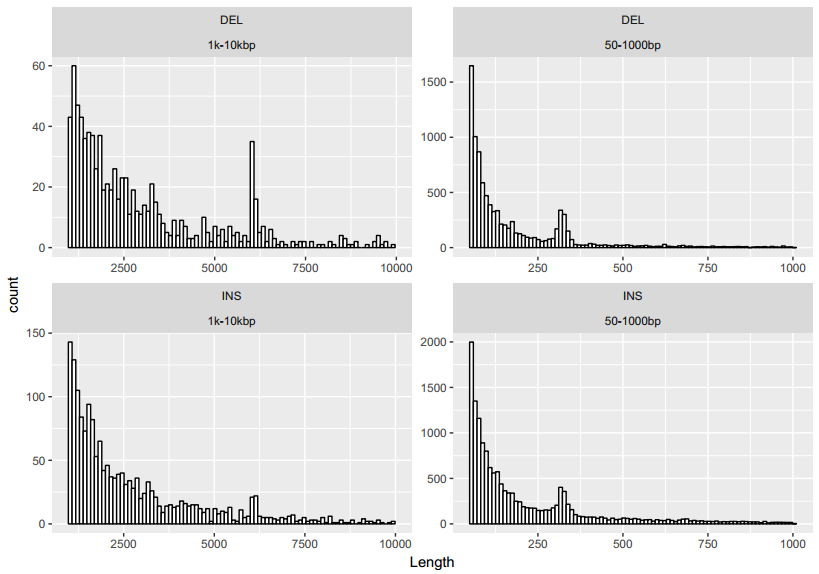

#### Figure S10 Size distributions for 0~1000bp and 1000~10000bp ranges for candidate SV calls from PacBio CCS assembly. Alu and LINE insertion/deletion events are at around 300 bp and 6000 bp.

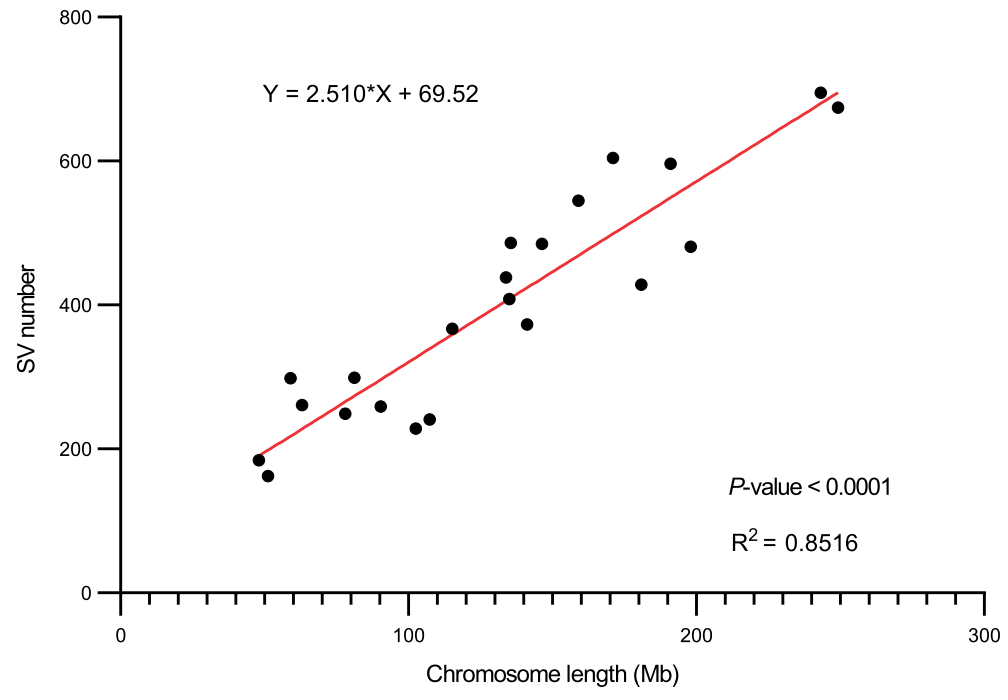

#### Figure S11 The relationship between chromosome length and the number of SVs

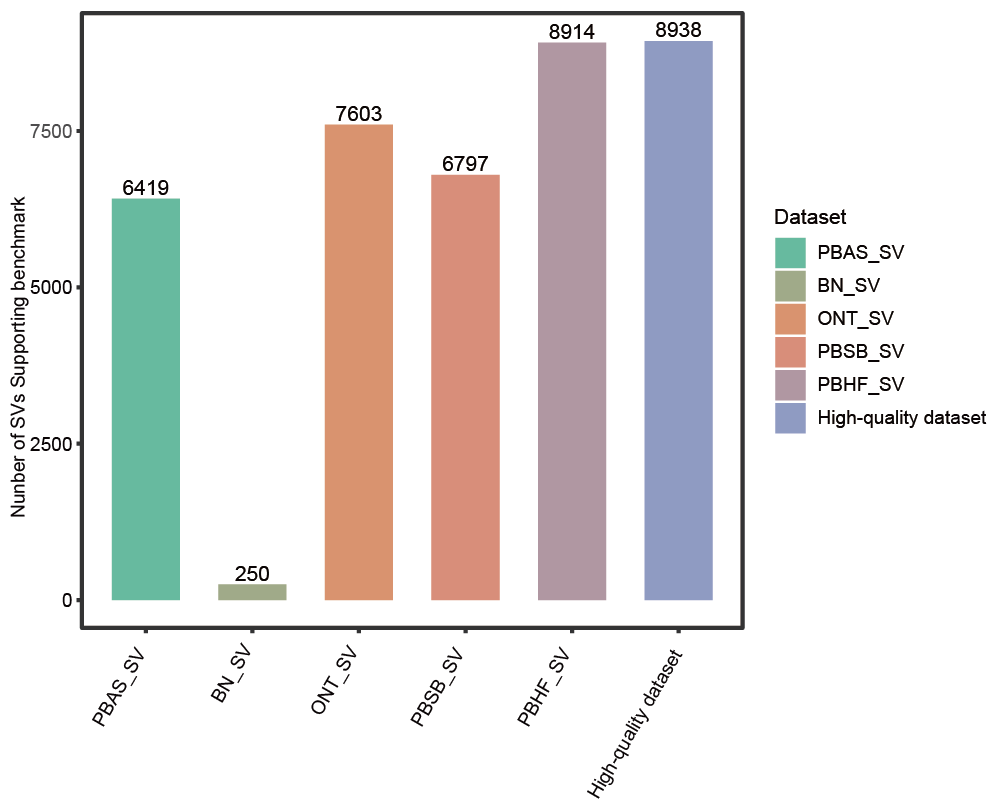

#### Figure S12 The number of unique SVs and SVs overlapping among different candidate SV callsets

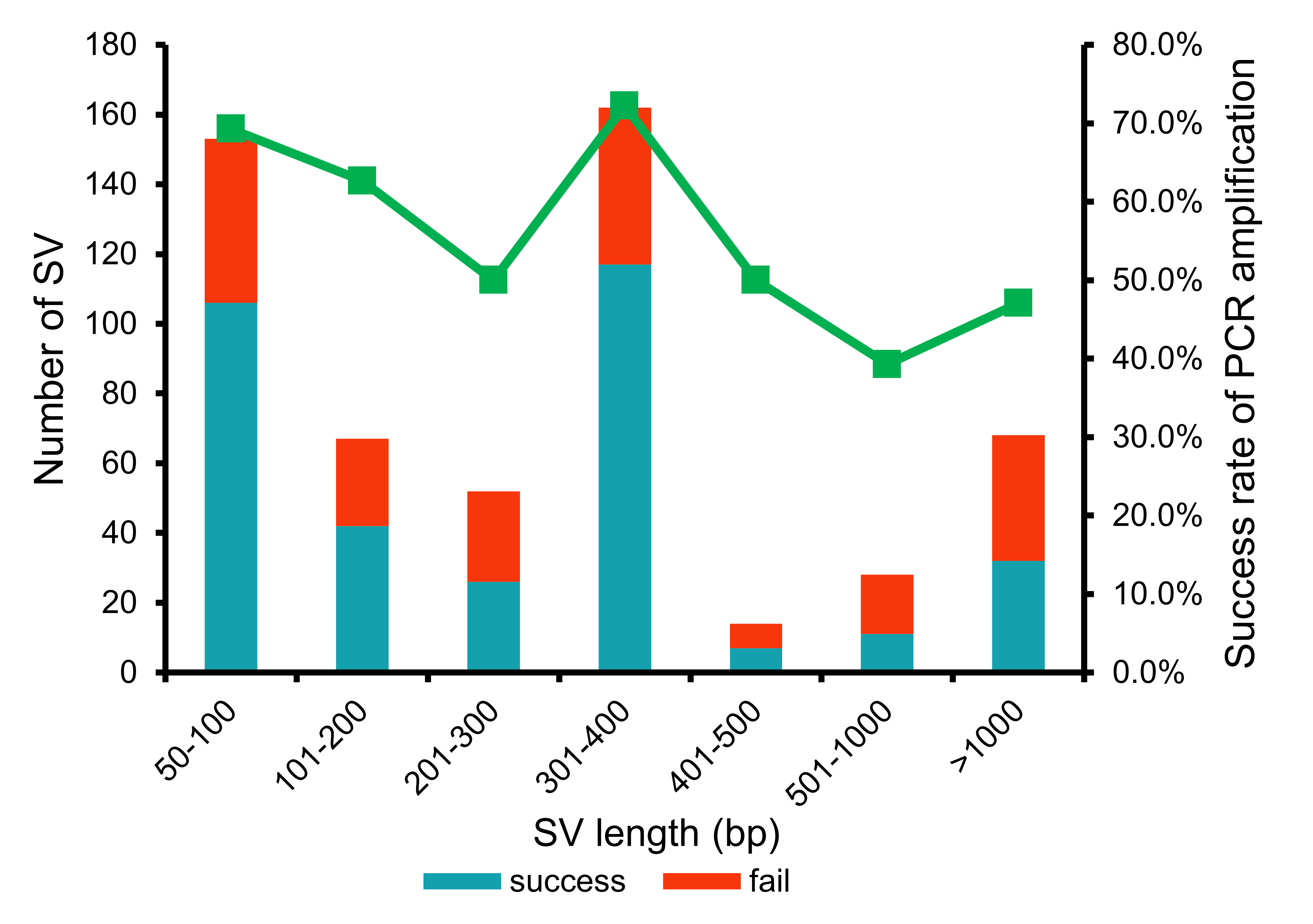

#### Figure S13 Distribution of PCR Amplification results in different length ranges

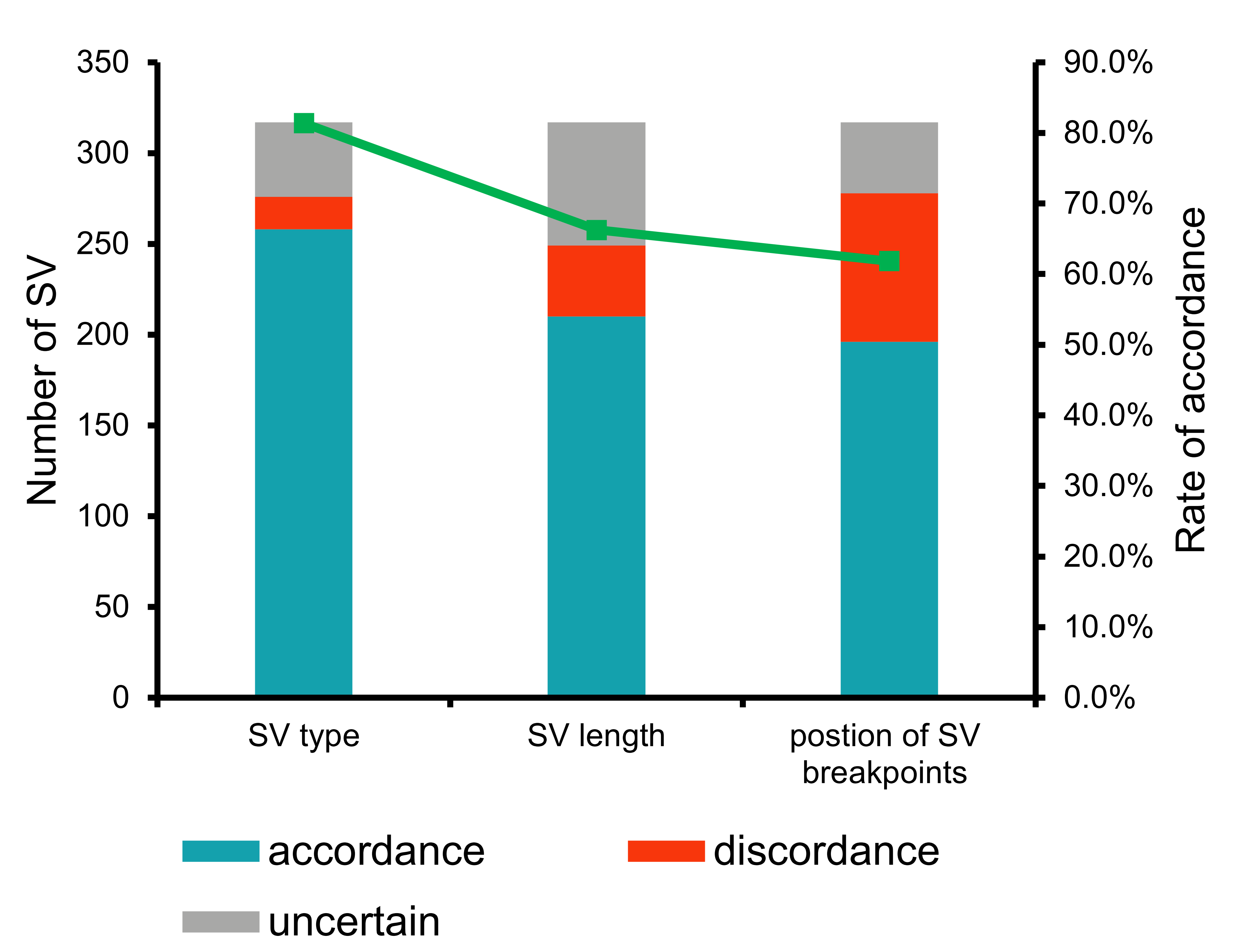

#### Figure S14 Consistency rates of SV type, SV length, and breakpoint position between Sanger sequenced variants and high-confidence SV calls with uncertain sites not excluded

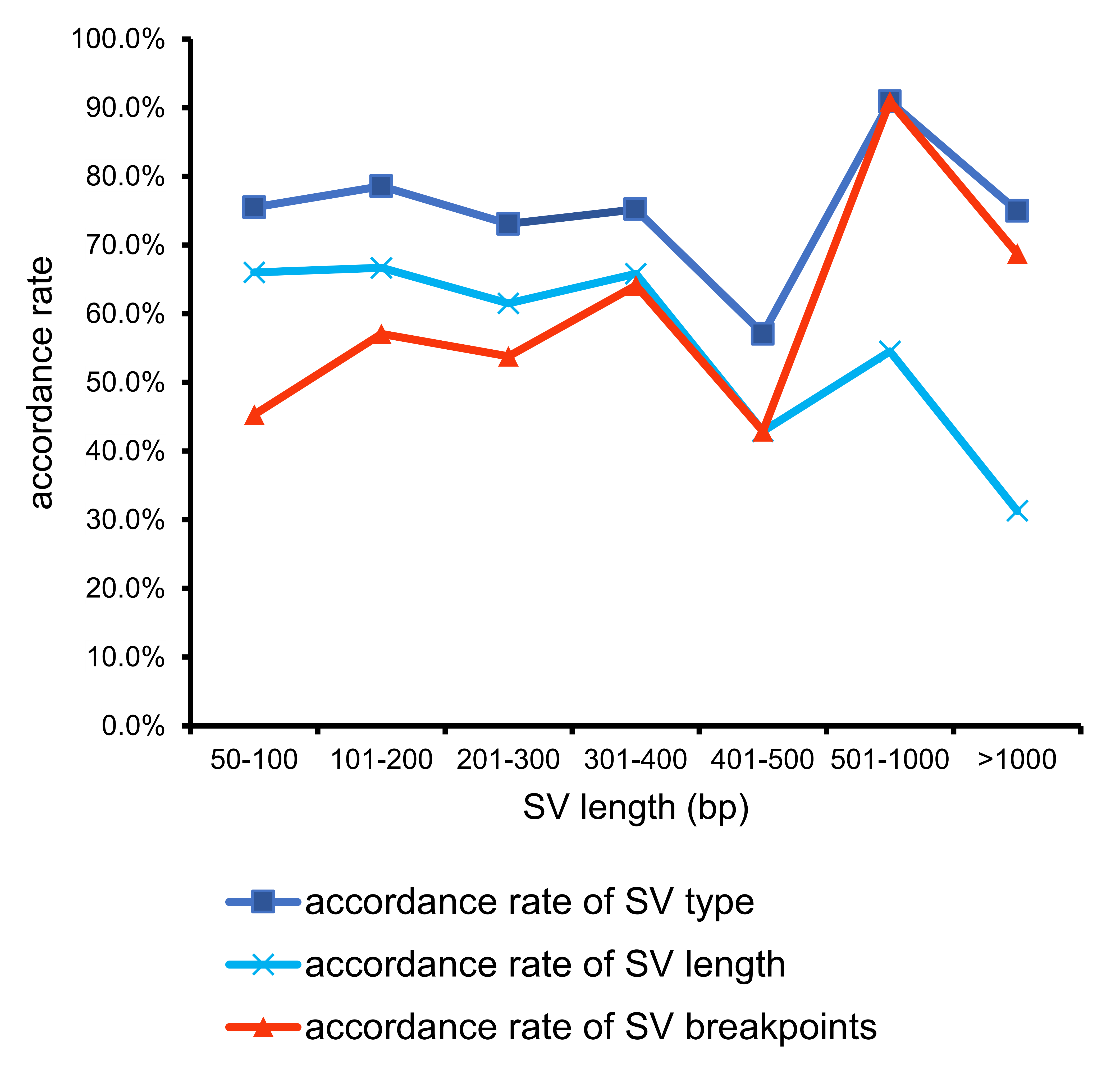

#### Figure S15 Consistency rates of SV type, SV length, and SV position between Sanger sequenced variants and high-confidence SV calls in different length ranges. Uncertain sites were not excluded.

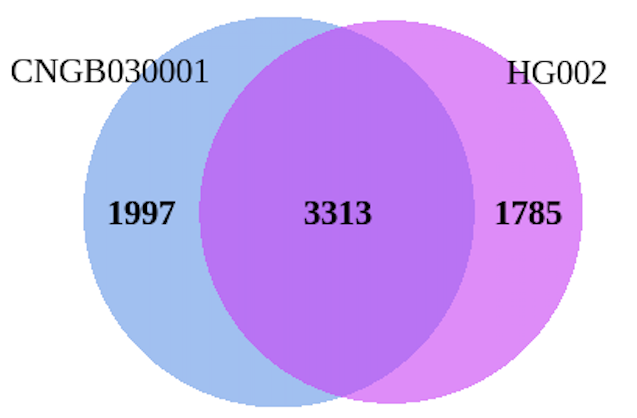

**Figure S16** Comparison of the Asian CNGB030001 benchmark and the GIAB HG002 benchmark within the 1.33 Gb overlapping benchmark regions.

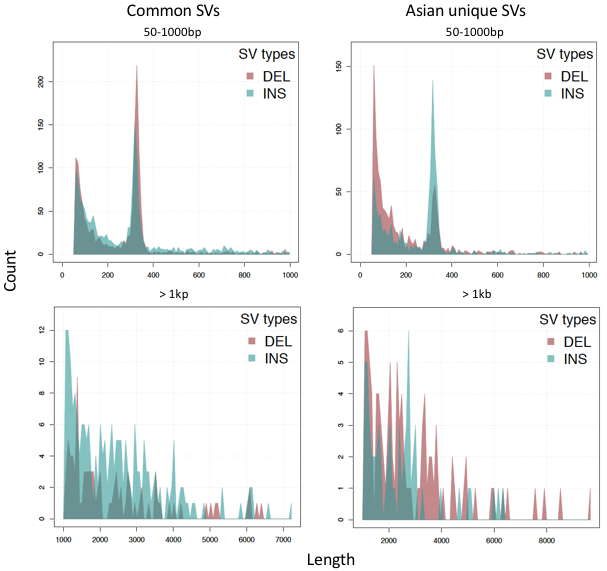

#### Figure S17 The size distributions for 50~1000bp and > 1kb ranges for common and Asian unique SVs by comparing GIAB benchmark to the Asian CNGB030001 benchmark.

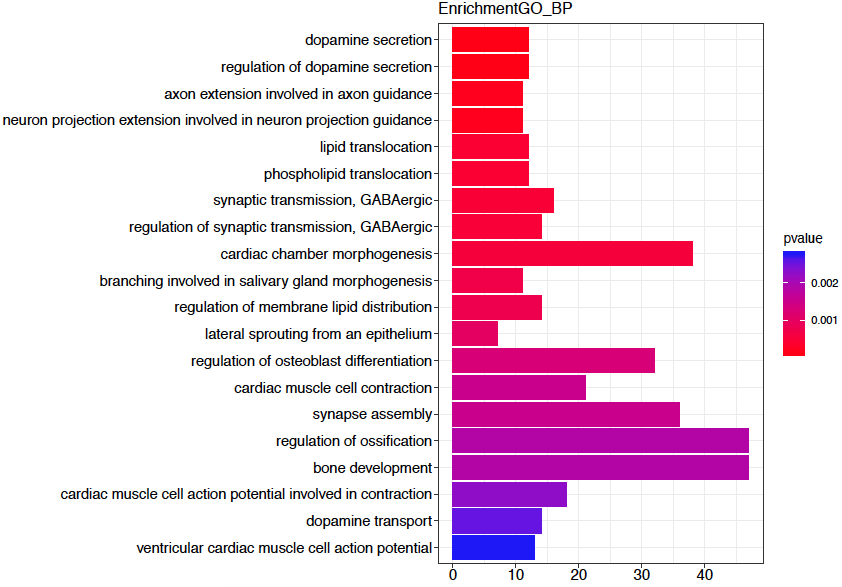

**Figure S18** GO enrichment of Asian-benchmark-specific SVs.

### Supplementary Tables

#### Table S1 Counts of identified high-confidence SVs on each chromosome

| Chromosome | DEL number | INS number | Total number |
| --- | --- | --- | --- |
| 1 | 318 | 356 | 674 |
| 2 | 342 | 353 | 695 |
| 3 | 234 | 247 | 481 |
| 4 | 333 | 263 | 596 |
| 5 | 230 | 198 | 428 |
| 6 | 297 | 307 | 604 |
| 7 | 281 | 264 | 545 |
| 8 | 244 | 241 | 485 |
| 9 | 169 | 204 | 373 |
| 10 | 257 | 229 | 486 |
| 11 | 178 | 230 | 408 |
| 12 | 207 | 231 | 438 |
| 13 | 169 | 198 | 367 |
| 14 | 120 | 121 | 241 |
| 15 | 99 | 129 | 228 |
| 16 | 129 | 130 | 259 |
| 17 | 148 | 151 | 299 |
| 18 | 130 | 119 | 249 |
| 19 | 148 | 150 | 298 |
| 20 | 121 | 140 | 261 |
| 21 | 83 | 101 | 184 |
| 22 | 75 | 87 | 162 |
| X | 75 | 91 | 166 |
| Y | 5 | 6 | 11 |
| Total | 4,387 | 4,540 | 8,938 |

#### Table S2 Trio binning of CCS reads

| *k*-mer | Parental CCS reads | Maternal CCS reads | Unknown | hapA/hapB | Assigned percentage (%) |
| --- | --- | --- | --- | --- | --- |
| 21 | 1,871,748 | 1,925,096 | 1,774,456 | 1.03 | 68.15 |
| 41 | 1,809,763 | 2,029,544 | 1,731,993 | 1.12 | 68.91 |
| 51 | 1,876,119 | 2,186,313 | 1,508,868 | 1.17 | 72.92 |
| 61 | 1,936,408 | 2,440,087 | 1,194,805 | 1.26 | 78.55 |
| 81 | 830,662 | 1,851,591 | 2,889,047 | 2.23 | 48.14 |
| Join | 1,941,539 | 2,374,195 | 1,255,566 | 1.22 | 77.46 |

#### Table S3 Counts of overlapping SVs from NGS platforms in comparison to the SV benchmark

|  | INS | | | | DEL | | | |
| --- | --- | --- | --- | --- | --- | --- | --- | --- |
|  | MGISEQ  2000 | % | NovaSeq  6000 | % | MGISEQ  2000 | % | NovaSeq  6000 | % |
| Manta | 1,397 | 39.51% | 1,000 | 28.28% | 1,901 | 56.81% | 2,100 | 62.76% |
| GRIDSS | 236 | 6.67% | 505 | 14.28% | 1,372 | 41.00% | 1,672 | 49.97% |
| LUMPY | 0 | 0.00% | 0 | 0.00% | 1,282 | 38.31% | 1,035 | 30.93% |
| BreakDancer | 39 | 1.10% | 5 | 0.14% | 1,386 | 41.42% | 1,277 | 38.16% |
| SV Benchmark | 3,536 |  |  |  | 3,346 |  |  |  |
